# Cell-free multi-omics analysis reveals tumor status-informative signatures in gastrointestinal cancer patients’ plasma

**DOI:** 10.1101/2023.01.31.526431

**Authors:** Yuhuan Tao, Shaozhen Xing, Shuai Zuo, Pengfei Bao, Yunfan Jin, Yu Li, Yingchao Wu, Shanwen Chen, Xiaojuan Wang, Yumin Zhu, Ying Feng, Xiaohua Zhang, Xianbo Wang, Qiaoran Xi, Qian Lu, Pengyuan Wang, Zhi John Lu

**Affiliations:** MOE Key Laboratory of Bioinformatics, Center for Synthetic and Systems Biology, School of Life Sciences, Tsinghua University, Beijing 100084, China; Institute for Precision Medicine, Tsinghua University, Beijing 100084, China; Translational Cancer Research Center, Division of General Surgery, Peking University First Hospital, Beijing 100034, China; Hepatopancreatobiliary Center, Beijing Tsinghua Changgung Hospital, Tsinghua University, No.168, Litang Road, Changping District, Beijing 102218, China; MOE Key Laboratory of Population Health Across Life Cycle, NHC Key Laboratory of Study on Abnormal Gametes and Reproductive Tract, Anhui Provincial Key Laboratory of Population Health and Aristogenics, Department of Maternal, Child and Adolescent Health, School of Public Health, Anhui Medical University, Hefei 230032, Anhui, China; Department of Integrative Medicine, Beijing Ditan Hospital, Capital Medical University, Beijing 100015, China

**Keywords:** Multi-omics, Cell-free DNA, Cell-free RNA, Cancer diagnosis, Liquid Biopsy

## Abstract

During cancer development, host’s tumorigenesis and immune signals are released to and informed by circulating molecules, like cell-free DNA (cfDNA) and RNA (cfRNA) in blood. However, these two kinds of molecules are still not systematically compared in gastrointestinal cancer. Here, we profiled 4 types of cell-free omics data from colorectal and stomach cancer patients, and assayed 15 types of genomic, epi-genomic, and transcriptomic variations. First, we demonstrated that the multi-omics data were more capable of detecting cancer genes than the single-omics data, where cfRNAs were more sensitive and informative than cfDNAs in terms of detection ratio, variation type, altered number, and enriched functional pathway. Moreover, we revealed several peripheral immune signatures that were suppressed in cancer patients and originated from specific circulating and tumor-microenvironment cells. Particularly, we defined a γδ-T-cell score and a cancer-associated-fibroblast (CAF) score using the cfRNA-seq data of 143 cancer patients. They were informative of clinical status like cancer stage, tumor size, and survival. In summary, our work reveals the cell-free multi-molecular landscape of colorectal and stomach cancer, and provides a potential monitoring utility in blood for the personalized cancer treatment.

## Introduction

The extracellular nucleic acid molecules include cell-free DNA (cfDNA) and cell-free RNA (cfRNA). They are usually fragmented but not fully degraded in plasma, due to protection of extracellular vesicle (EV), or binding protein like nucleosome for cfDNA and ribonucleoprotein (RNP) for cfRNA. These extracellular molecules have been widely used in cancer diagnosis and prognosis, because cancer alterations of tumor cells can be detected from cfDNAs and cfRNAs in the circulating blood(1). In addition, tumor microenvironment and peripheral immune system are also crucial in characterizing a patient’s status during cancer treatment. For instance, a patient’s stromal cell activity and systematic immune response will affect his clinical treatment outcome (e.g., immunotherapy)(2, 3). Cell-free molecules, especially cfRNAs, contain signals released from these non-tumor cells as well(4).

Many cfDNA features, such as methylation, mutation, copy number, fragment pattern, and nucleosome footprint, have been utilized for noninvasive diagnosis and prognosis of diseases(5–7). Meanwhile, many cfRNA features can also be used as biomarkers, such as the abundance of miRNAs(8) and circRNAs(9), fragment copy and alternative splicing of mRNAs and lncRNAs(10–12). Moreover, many other RNA regulation events altered in tumor cells, such as RNA editing(13), can potentially be utilized in liquid biopsy as well.

In tumor cells and tissues, many studies have demonstrated that multi-omics data provided a more comprehensive understanding of diseases than single-omics data(14, 15). In liquid biopsy, integrating multiple cell-free molecules also enhanced the diagnosis power. For instance, cfRNA and cfDNA were used together to detect EGFR mutation in plasma(16); 61 DNA mutations and 8 proteins were combined in a multi-analyte blood test for cancer(17). However, cell-free multi-omics data of cfDNAs and cfRNAs have not been systematically investigated in cancer, such as colon cancer (CRC) and stomach cancer (STAD), two of the most common types of gastrointestinal cancer. Existing methods for diagnosis and treatment monitoring of these two cancers, such as endoscopy and tissue biopsy, still lack convenience, sensitivity, and information of molecular mechanisms(18). Therefore, uncovering the shared and distinct cell-free signatures of these two gastrointestinal cancers will help us understand their extracellular biology, and provide noninvasive monitoring utilities.

Here we present a systematic evaluation of cell-free multi-omics data, including methylated cfDNA immunoprecipitation sequencing (cfMeDIP-seq), cfDNA whole genome sequencing (cfWGS), total and small cfRNA sequencing (cfRNA-seq) data. Each group of the matched multi-omics data was sequenced from 2-3 mL plasma sample. Using colorectal cancer and stomach cancer as two example gastrointestinal cancer types, we investigated multiple alterations of cfDNAs and cfRNAs, providing a cell-free multi-molecular landscape.

## Results

### Profiling cell-free multi-omics and data quality control

To study cell-free multi-omics and compare them in colorectal cancer and stomach cancer, we sequenced 4-omics data for 161 individuals (Fig. 1a, see details in Methods, Extended Data Fig. 1a-c, Supplementary Table 1). The data’s qualities were well controlled (see quality control steps in Methods, Supplementary Tables 2,3): intra-omics correlation between samples was above 0.75 in every single omics; inter-omics correlations were mostly close to zero (Extended Data Fig. 1d); concentration, read length, and read distribution of the data were consistent with previous studies(5, 12, 19) (Extended Data Fig. 2). As expected, the sequenced reads’ distributions of cfDNAs and cfRNAs were very different: cfRNA-seq provided abundant information in exonic regions, while cfDNA-seq provided wide information in all exonic, intronic and intergenic regions (Fig. 1b).

**Figure 1.**
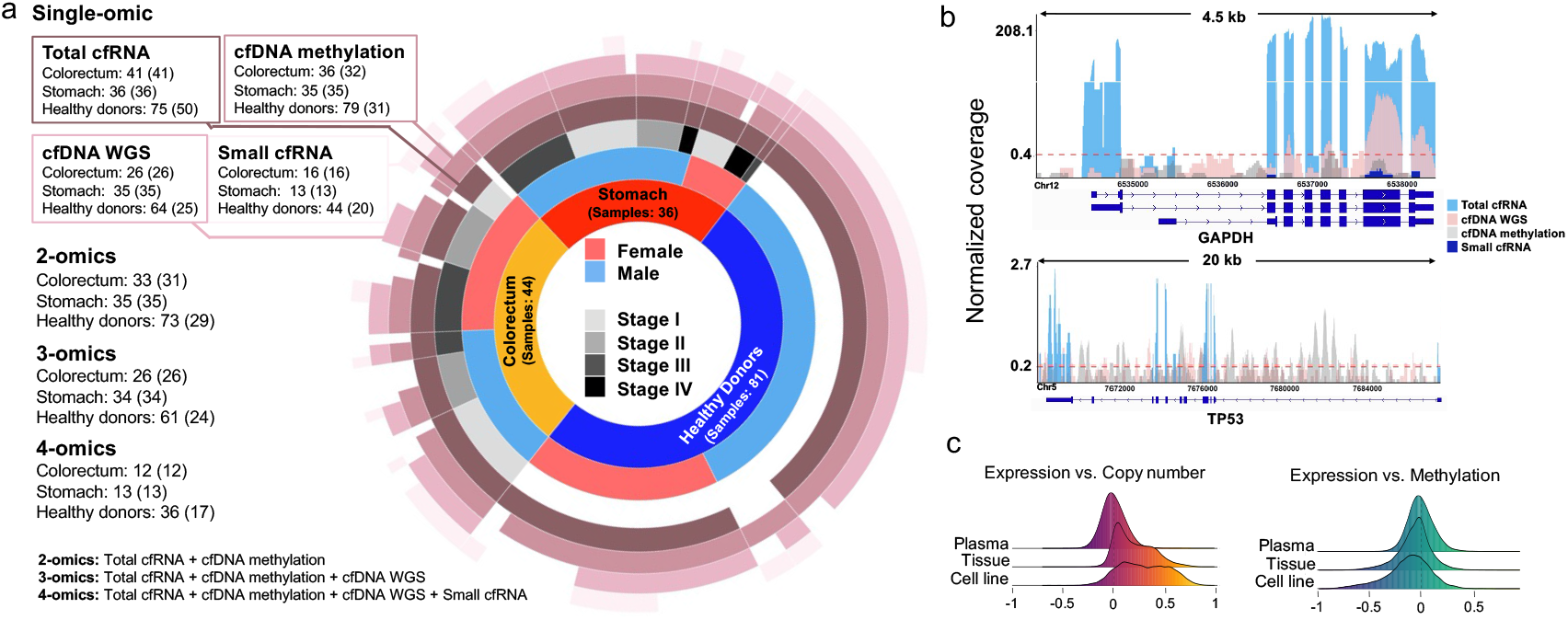
Cell-free multi-omics data summary and quality control. a, Cell-free multi-omics data in plasma. Nurnbers inuide and outside brackets are datasets and samples, respectively, where some samples were mixed for sequencing. The gap in the ring means no paired data. b, Multi-omics reads mapped on a housekeeping gene, *GAPDH*, and a tumor suppressor gene, *TP53*. The coverage is normalized by total mapped reads. Red dashed line: the average coverage of cfDNA reads mapped on a gene. c, Density plots of multi-omics correlation coefficients of genes in tissues (TCGA), cell lines (CCLE), and plasma (this study).

We compared correlations between multi-omics in plasma (our data), cell lines (CCLE data(20)), and tissues (TCGA data(21)) (Fig. 1c). Different from the data in cells, signals of cfDNAs and cfRNAs were not well correlated, probably due to the heterogenous origins of cfDNAs and cfRNAs(4, 22). From another perspective, combining orthogonal information of different cell-free molecules could potentially detect cancer with better capacity than using single-omics data in plasma. Thus, we combined the multi-omics data to detect cancer genes and compared their detection capacities in the following analyses.

### Combination of cell-free multi-omics data enhanced the detection of cancer genes in plasma

We comprehensively profiled and calculated 15 cell-free molecular variation events using our multi-omics data (see Methods). These variations can be used to examine cancer patients and healthy donors (HDs) in a multidimensional view of individual samples (Fig. 2a) and genes (Fig. 2b). For instance, *TP53* in tumors is usually depleted at DNA copy number level and downregulated at RNA expression level in cancer patients (Fig. 2b). We found that certain cancer patients were not able to be simultaneously detected by both cfDNA and cfRNA at 95% specificity, suggesting that combination of cfDNA and cfRNA data would improve detection capacity of *TP53*. The 95% specificity was defined by variation values in healthy donors (HDs): an individual with a variation value (e.g., mutation ratio or abundance level) above 95% quantile of HDs was identified as an outlier.

**Figure 2.**
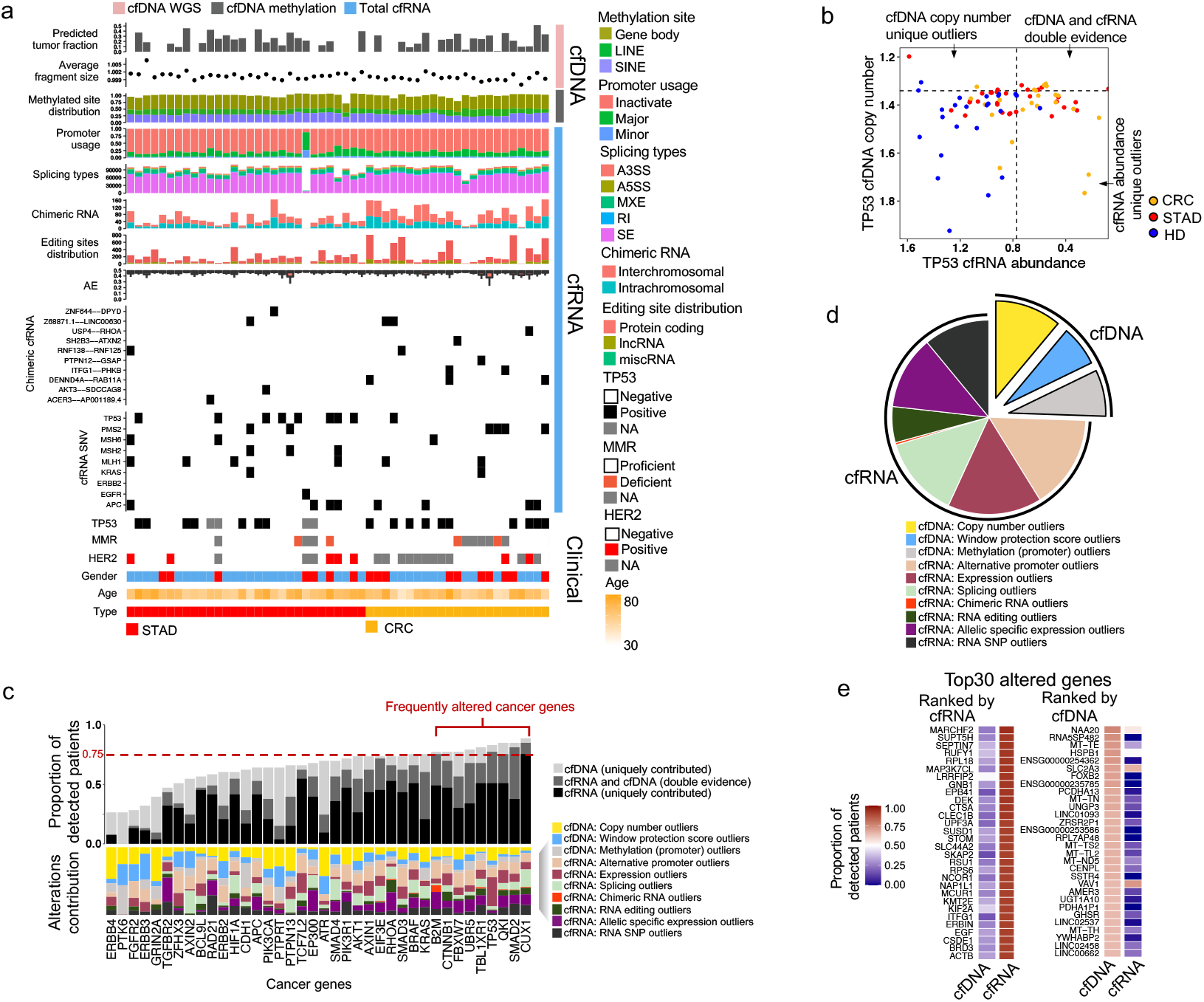
Detection of the cancer genes using different types of variations. **a,** Overview of the plasma multi-omics variation atlas. **b,** cfDNA copy number and cfRNA abundance for *TP53*. HD: healthy donor, CRC: colorectal cancer, STAD: stomach cancer. The dot lines represent 95% specificity defined by HDs. **c,** Detection capacity for each cancer gene combing different variations derived from cfDNA (cfWGS and cfMeDIP-seq) and total cfRNA-seq data. Frequently altered genes are defined by >75% detection ratio. **d,** Distribution of variation types among all genes that are frequently altered. **e,** Altered genes with top detection ratios ranked by cfRNA and cfDNA, respectively.

We quantified the detection capacity (sensitivity) of a cancer gene as the proportion of cancer patients being detected at 95% specificity, then investigated each variation type for a set of pre-defined cancer genes. This cancer gene set was defined by referring to the COSMIC hallmark cancer genes(23), where 38 genes with somatic mutations annotated as *colorectal* or *gastric* were used. Our data demonstrated that multiple variations’ combination leads to a great increase in detection capacity, while a single alteration usually detected less than half of the patients (Fig. 2c). Alterations of 9 cancer genes, like *CUX1, SMAD2, QKI*, and *TP53*, were detected at cfDNA or cfRNA level in most patients (>75%), suggesting that these genes were frequently altered in the cancer patients. Notably, certain cfRNA variations were the major contributors, for instance, RNA alternative promoter (average ratio:17.7%), RNA expression (average ratio: 13.3%), allele specific expression (average ratio: 12.5%), and RNA splicing (average ratio: 11.4%).

In summary, our result demonstrated that combining cell-free multi-omics data enhanced the detection capacity (i.e., sensitivity score at 95% specificity) for each of the predefined cancer genes, where cfRNA variations often contributed more to the sensitivity than cfDNA variations.

### cfRNA variations were more sensitive in cancer-related gene detection than cfDNA variations

We further expanded the investigation from the pre-defined cancer genes to the genes altered in cancer patients (i.e., cancer-related genes). A gene was defined as a frequently altered one if found as an outlier (beyond 95% of HDs) in more than 75% of the cancer patients. Consistent with the pre-defined cancer genes, the frequently altered genes were mostly identified by cfRNA variations (Fig. 2d). In other words, the cfRNA variations tend to be more sensitive than cfDNA in cancer. This finding in plasma is similar to a multi-omics study in tumor cells, which revealed that RNA variations accounted for 78.23% of all identified alterations in 731 genes with significant recurrent aberrations(14).

We also listed 30 genes with top sensitivity (proportion of detected patients) altered at cfDNA or cfRNA level (Fig. 2e). The data further demonstrated that cfRNA variations were more sensitive than cfDNA variations: top genes altered at cfDNA level usually had lower sensitivity than those top ones at cfRNA level. Moreover, we also found that the top genes found by cfDNA and cfRNA were very different, indicating that their information was complementary to each other in plasma. For instance, many top genes (e.g., *MAP3K7CL, DEK, CLEC1B*, and *SKAP2*) altered at cfRNA level are functionally related to oncogenes and immune pathways; among the top genes altered at cfDNA level, mitochondrial DNA (mtDNA) frequently occurred, which could be related to the increased metabolism in cancer(24).

### Identification of various cell-free molecules’ differential alterations in cancer patients

In addition to analyzing the variations altered in the individual patients, we used a statistical differential analysis to identify differentially altered variations between two groups, e.g., cancer patients and HDs, colorectal and stomach cancer patients (see Methods, Fig. 3a). Among these differential alterations, we found that cfRNA abundance, cfDNA methylation level, and cfDNA window protection score (WPS) were mostly altered in stomach cancer, while cfRNA abundance, cfRNA SNV, and cfDNA WPS were mostly altered in colorectal cancer, in terms of differential number (Fig. 3a). Many well-known cancer alterations were identified from the differential analysis. For instance, we found that *KRAS’s* cfRNA abundance was significantly higher in cancer patients (*edgeR* exactTest*, P*-value = 0.011). Moreover, *KRAS’s* promoter region was also more open in cancer patients according to its cfDNA WPS (Fig. 3b). Another example is the cfDNA methylation level of *PGRMC1*, which is a carbon-monoxide-sensitive molecular switch associated with *EGFR(25*). We found that the cfDNA of *PGRMC1* was hypo-methylated at its promoter region in the cancer patients (*edgeR* exactTest, *P*-value = 0.047) (Fig. 3b).

**Figure 3.**
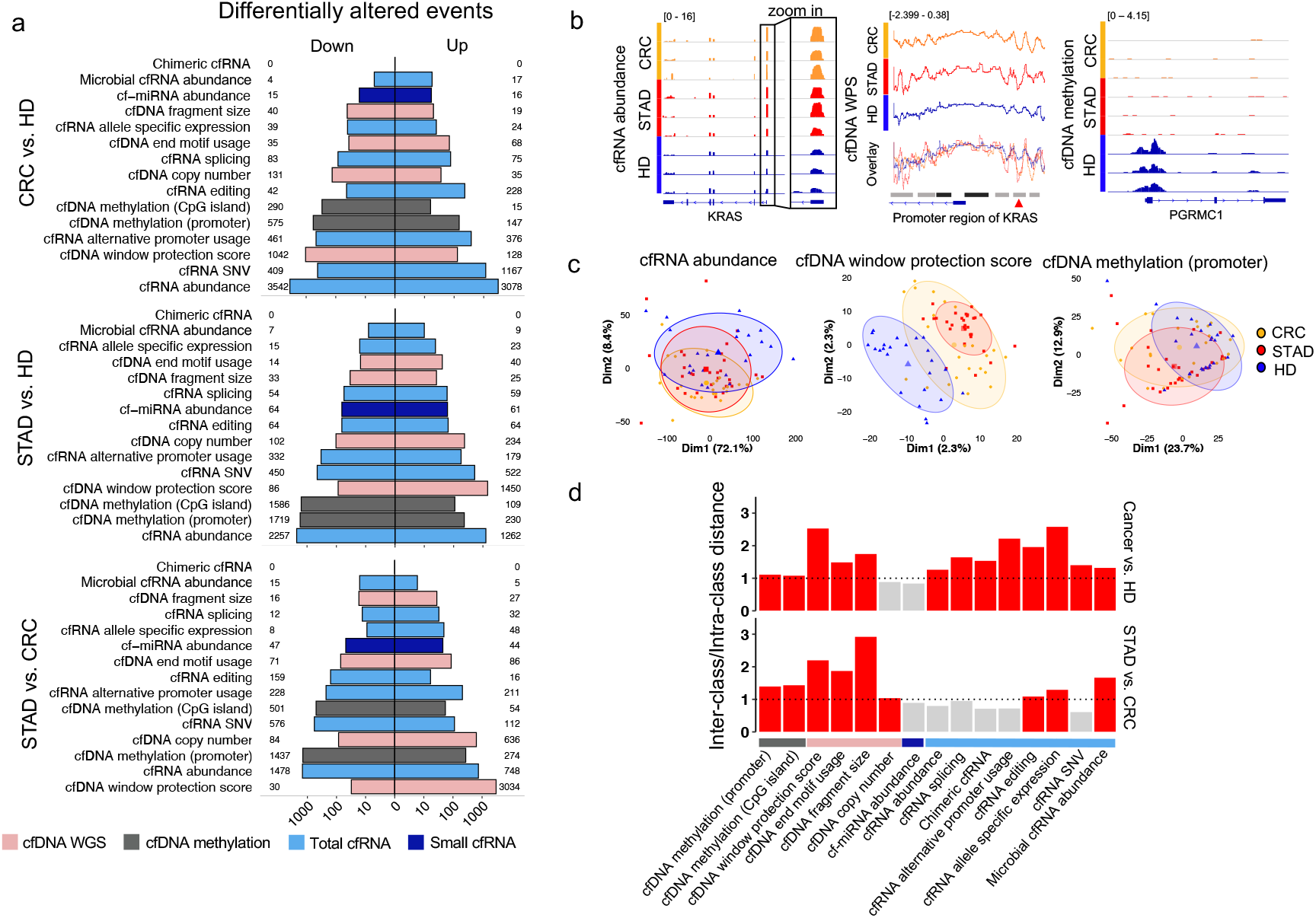
Various cell-free molecules’ differential alterations. **a**, Numbers of the differentially altered events in plasma between cancer patients and healthy donors (HDs), and between colorectal (CRC) and stomach cancer (STAD). **b**, Examples of *KRAS’s* cfRNA abundance, *KRAS’s* window protection score (WPS), and *PGRMC1’s* cfDNA methylation. The blue blocks, lines and arrows below each panel are gene models. Black blocks above promoter region of *KRAS:* promoter regions; grey blocks: enhancer regions; red arrow: open regions in cancer. **c**, PCAs of 3 representative differential alterations among cancer patients and HDs. **d**, Ratio of inter-class distance over intra-class distance for each type of differential alteration. Ratios larger than 1 (dashed line) are colored red.

We examined different alterations’ capacities in cancer classification (Fig. 3c, Extended Data Fig. 3). Ratio of inter-class distance over intra-class distance was used to quantify the classification capacity for each type of differential alternations (Fig. 3d). We found that the alterations derived from the cfDNA WGS data performed well in both cancer detection (i.e., cancer patients vs. healthy donors) and cancer type classification (i.e., colorectal vs. stomach cancer). The cfMeDIP-seq data worked slightly better in cancer-type classification than in cancer detection. Meanwhile, miRNA abundance derived from the small cfRNA-seq data performed not well in either case, while the alterations derived from the total cfRNA-seq data usually worked better in cancer detection than in cancer type classification. In addition, microbial cfRNA abundance derived from the total cfRNA-seq data better classified the two cancer types than the features of human cfRNAs (Fig. 3d), which is consistent with the result we previously reported(12).

### Suppressed immune signatures in plasma revealed by the cell-free multi-omics data

We assayed the enriched pathways for the differential alterations (Fig. 3a) between cancer patients and HDs, and found that cfRNA data were relatively more informative than cfDNA data in terms of the number (Fig. 4a) and function (Fig. 4b) of the enriched pathways. In the up-regulated genes, we revealed cancer-related pathways significantly enriched in the alterations detected by cfDNA copy number and cfRNA abundance (*edgeR* exactTest, *P*-value < 0.05). In addition, we found several immune pathways (e.g., T cell and B cell receptor signaling pathways) enriched in the genes with down-regulated cfRNA abundance in the patients (Fig. 4b, Extended Data Fig. 4), suggesting an immunosuppression state of these pathways in the cancer patients’ plasma. For instance, *CD8A*, an indicator of cytotoxic T cells, and *ZAP-70*, a critical gene in activating downstream signal transduction pathways in T cells(26), were both down-regulated in the cancer patients’ plasma. *CD19*, an indicator of B cells was also found to be down-regulated in the patients. Particularly, *PD-L1*, a well-known immune suppressor in cancer, was significantly up-regulated in the colorectal cancer patients’ plasma (Fig. 4c).

**Figure 4.**
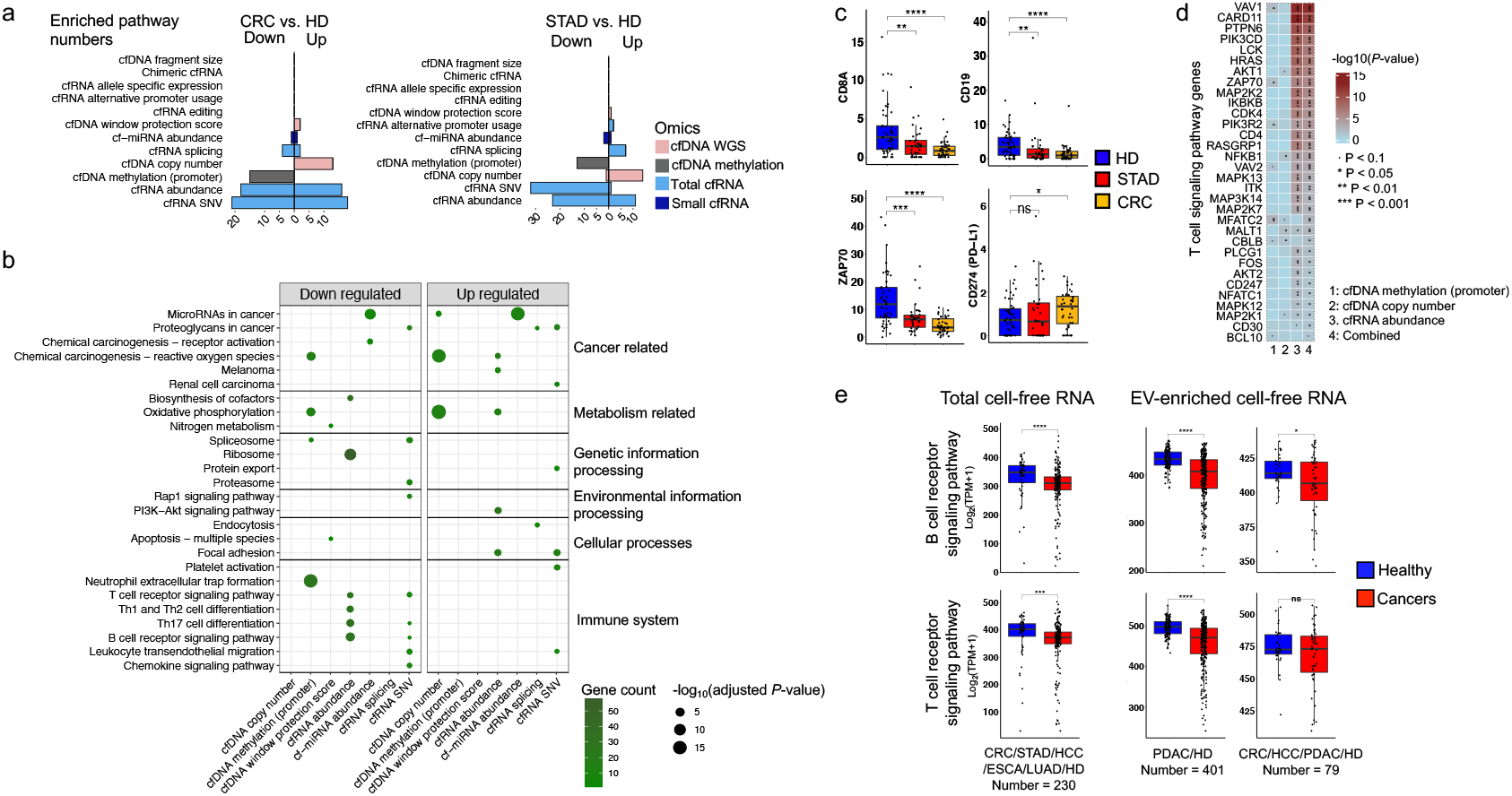
Enriched fu net io na l pathwa ys of the differential alterations. **a,** Numbers of the enriched pathways for the differentially altered genes defined by different variation events in cancer patients’ plasma. **b,** KEGG terms of the top enriched pathways for each differential alteration. **c,** Differential cfRNA abundance for the example genes in the immune pathways. Y axis is TPM. **d,** Example genes in T cell receptor signaling pathway altered at different omics levels. *P*-value represents significance of the differential alterations. Combined *P*-value was calculated by *Activepathways*. **e,** Down-regulated T cell and B cell receptor signaling pathways calculated by the public total and EV-enriched cfRNA-seq datasets. Single-tailed Wilcoxon rank-sum test was used. *P < 0.05, **P < 0.01, ***P < 0.001, ****P < 0.0001, ns: not significant. HD: healthy donor, CRC: colorectal cancer, ESCA: Esophageal carcinoma, HCC: hepatocellular carcinoma, LUAD: lung adenocarcinoma, PDAC: pancreatic ductal adenocarcinoma, STAD: stomach cancer.

Multi-omics pathway enrichment (see Methods) also confirmed this immunosuppression, while most of the down-regulated events were found by cfRNA rather than cfDNA (Fig. 4d, Extended Data Fig. 5). For instance, among the differentially altered genes enriched in the T cell receptor signaling pathway, 28 out of the 32 genes were significantly changed with cfRNA abundance, but only 7 genes were changed with cfDNA alterations.

Subsequently, we validated the immunosuppression state in 710 published cell-free data, including total cfRNA and EV-enriched cfRNA data of colorectal cancer (CRC), stomach cancer (STAD), esophageal cancer (ESCA), hepatocellular carcinoma (HCC), Pancreatic ductal adenocarcinoma (PDAC), and lung adenocarcinoma (LUAD) (Supplementary Table 4). We found that the T cell and B cell receptor signaling pathways were also significantly down-regulated in the plasma of these cancer patients (Fig. 4e).

In summary, the cell-free multi-omics data uncovered many cancer-related functional pathways and signatures in plasma. Particularly, we revealed several down-regulated immune signatures in the cancer patient’s plasma, which were mostly contributed by cfRNAs.

### cfRNA features originated from specific circulating and tumor microenvironment cells

In order to trace the origins of these signatures derived from cfRNAs, we sequenced total RNAs in the paired samples of plasma, primary tumor, normal tissue adjacent tumor (NAT), and peripheral blood mononuclear cells (PBMCs) from 16 colorectal cancer patients and 6 healthy donors (Fig. 5a). We used a computational deconvolution method, EPIC(27), to estimate the origins/components of RNA-seq reads in plasma, tissue, and PBMCs. Remarkably, we found that the cfRNAs in plasma originated not only from blood cells like lymphoid and macrophages but also from tumor-microenvironment cells like endothelial cells and cancer-associated fibroblasts (CAFs). Actually, the cfRNAs in plasma captured a CAF signal that was hardly detectable by the PBMC RNAs (Fig. 5b). In summary, the data demonstrate that plasma cfRNAs contain signals not only from peripheral blood cells but from tissue cells.

**Figure 5.**
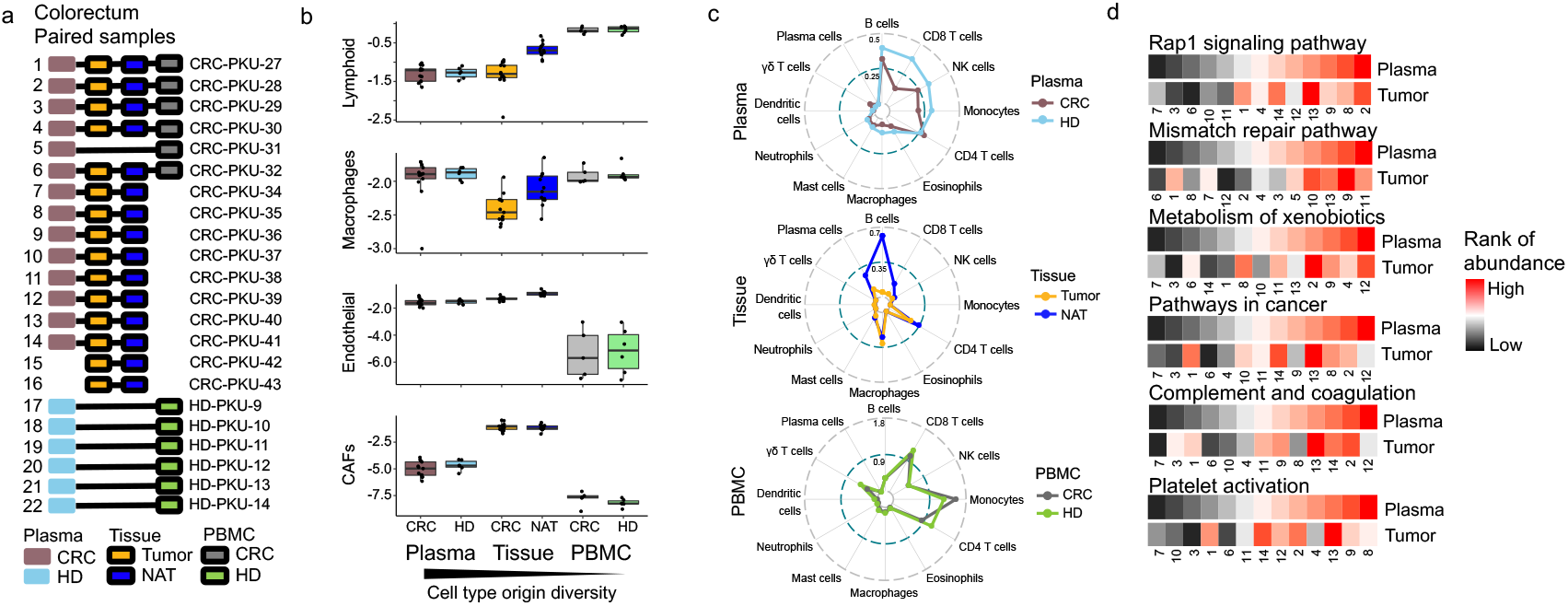
RNA expression signals compared in plasma, PBMC, and tumor. **a**, RNA sequencing data in the paired samples of colorectal cancer (CRC) patients. NAT: normal tissue adjacent tumor. **b**, Inferred signals originated from different cell types for 3 types of RNA-seq data (plasma, PBMC, and tissue). Y-axis is log10 transformed cell type ratio estimated by EPIC. **c**, Inferred relative abundance of more cell types. **d**, Correlated pathways of plasma and paired tumor samples. The abundance value of a pathway was averaged from the genes in this pathway. The numbers on the x-axis correspond to the sample identifiers in a.

In order to find out which type of cells were down-regulated in the cancer patients, we inferred immune cell abundance using the cfRNA-seq data based on the LM22 immune cell markers(28) (Fig. 5c). CD8 T cell abundance was substantially down-regulated in both cancer patients’ plasma (cancer patients vs. healthy donors, Wilcoxon rank sum test, *P*-value = 0.006) and primary tumor (primary tumor vs. NAT, Wilcoxon rank sum test, *P*-value = 0.009). In addition, other cell types with tumor-killing potential, such as B cells and NK cells, were also down-regulated in plasma and primary tumors. Notably, these down-regulated immune signatures detected in both plasma and tumors were not well detected by the PBMCs (Fig. 5c), indicating that plasma cfRNAs had the potential to better monitor cancer microenvironment and developing status than PBMC RNAs.

Furthermore, we calculated the gene expression correlation of more pathways between the paired tumor and plasma samples (Fig. 5d). Significant positive correlations between plasma and tumors were found in many pathways, such as Rap1 signaling pathway (Spearman correlation, R = 0.764, *P*-value = 0.002), mismatch repair (Spearman correlation, R = 0.698, *P*-value = 0.005), cancer-related pathway (Spearman correlation, R = 0.5, *P*- value=0.043), complement and coagulation cascades (Spearman correlation, R = 0.588, *P*- value = 0.019), platelet activation (Spearman correlation, R = 0.533, *P*-value = 0.032).

In summary, we have revealed positive correlations between plasma cfRNAs and tumor RNAs for certain cancer- and immune-related signatures. Plasma cfRNA’s capability of tracing tumor signals/origins suggests its utility as a noninvasive monitor of clinical status for cancer.

### Signatures derived from plasma cfRNAs informed clinical status of cancer patient

To prove that the signatures derived from the plasma cfRNAs are able to monitor clinical status of cancer patients, we assayed 143 total cfRNA-seq datasets from colorectal and stomach cancer patients, where previous data we published (GSE174302)(12) were also included. We calculated various cell type signature scores based on deconvolution of the total cfRNA-seq data (see Methods) (Fig. 6a, b). Remarkably, scores of specific cell types, such as γδ T cells, resting NK cells, M2 tumor-associated macrophages (TAM), and cancer-associated fibroblasts (CAFs), were highly correlated with the cancer stage status (Fig. 6c-f). For instance, the γδ-T-cell score was negatively correlated with cancer stage for colorectal and stomach cancer (Fig. 6a, c) and tumor size for colorectal cancer (Fig. 6g), consistent with its anti-tumor function(29, 30). In addition to the immune signatures, the cancer microenvironment score of CAFs was also positively correlated with the cancer stage (Fig. 6a, d). Consistently, when applying the plasma-derived CAF score to a TCGA cohort (1,006 colorectal and stomach cancer patients), we found that the CAF score was negatively correlated with patient’s survival (Fig. 6h).

**Figure 6.**
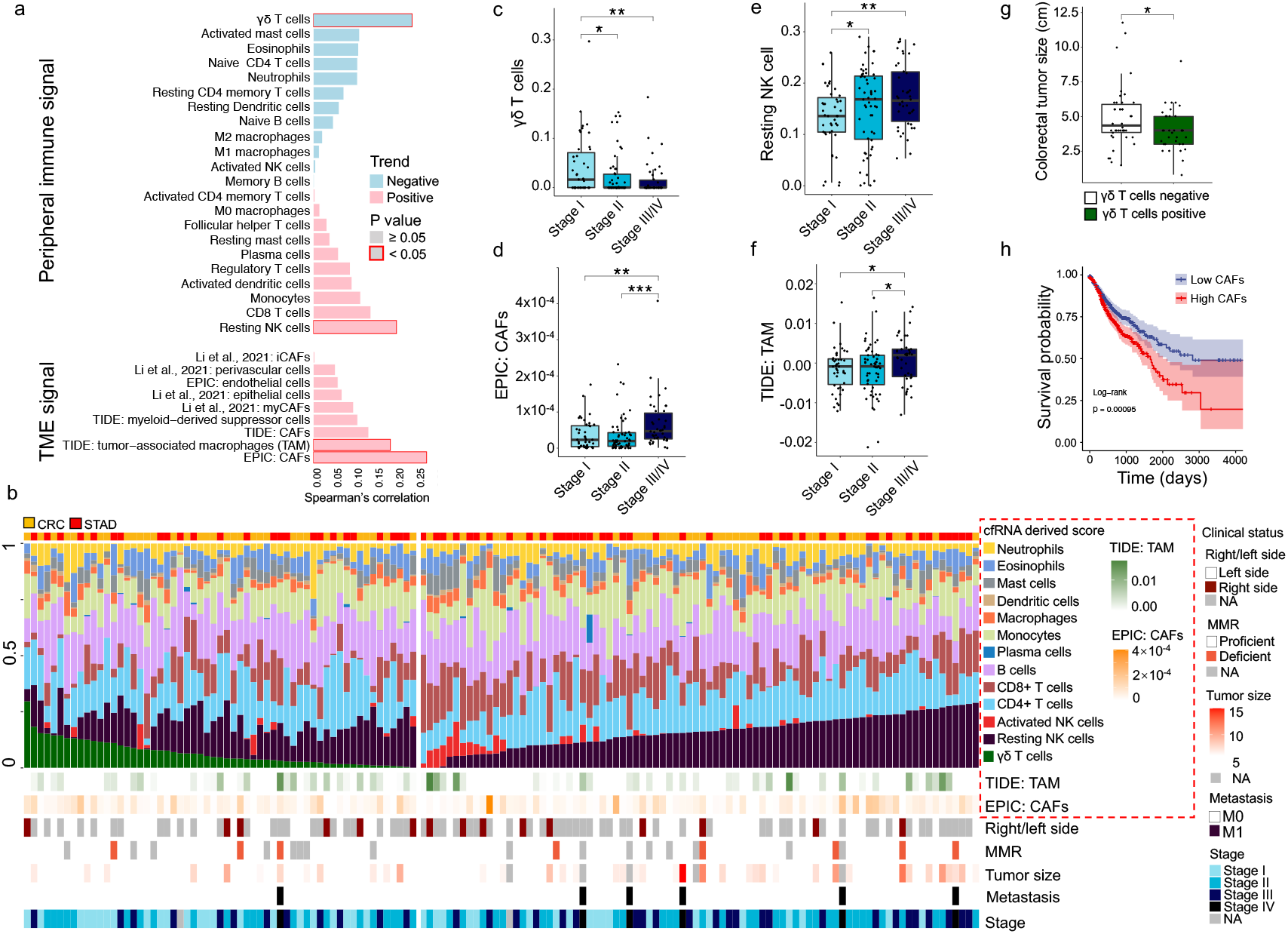
Clinical status-informative signatures derived from the plasma cfRNAs. **a,** Correlations between cancer stage and cfRNA-derived cell-type signatures for CRC and STAD patients. TME: tumor microenvironment, CAF: cancer-associated fibroblast, iCAF: inflammatory CAF, myCAF: myofibroblastic CAF. **b,** Overview of various cfRNA-derived signatures and individual clinical status, ranked by γδ-T-cell score and resting-NK-cell score. The cfRNA-derived scores of **c,**γδ T cells, **d,** EPIC: CAFs, **e,** Resting NK T cells, and **f,** TIDE: TAM, are shown in different cancer stages for CRC and STAD patients. **g,** Tumor sizes for 2 subtypes of CRC patients categorized by the γδ-T-cell scores (positive: >0; negative: =0) in plasma. Single-tailed Wilcoxon rank-sum tests were used. *P < 0.05, **P < 0.01, ***P < 0.001, ****P < 0.0001. **h,** Survival with high (top 50%) and low (bottom 50%) scores of EPIC: CAFs in the TCGA cohort of 475 COADs (colon cancer), 164 READs (rectal cancer), and 367 STADs (stomach cancer). Log-rank test was used for survival time comparison.

In summary, we have revealed specific immune (e.g., γδ T cells) and cancer-microenvironment (e.g., CAFs) signatures decomposed from plasma cfRNA-seq data, which can be used as predictive scores to monitor a patient’s clinical status like cancer stage, tumor size, and survival.

## Discussion

### Conclusion

In this study, we present a landscape of cell-free nucleic acids for colorectal and stomach cancer based on paired data of genome, epigenome, and transcriptome in plasma. Moreover, we have demonstrated the concept of multi-omics integration in liquid biopsy. Subsequently, we provide a cfRNA-based utility for the monitoring of cancer status.

### Clinical significance of monitoring cancer status using noninvasive biomarkers

Conveniently monitoring a patient’s status, like tumor size, cancer stage, and immune response, is very important during cancer treatment, such as immune therapy and neoadjuvant therapy(31). However, existing methods evaluating response and effectiveness of a treatment are still inconvenient and inaccurate(32). Liquid biopsy based on cfDNA/cfRNA biomarkers is a promising approach, because it is non-invasive, less painful, low cost, and convenient. The quantitative signatures/scores based on the noninvasive biomarkers would help doctors perform the best therapeutic regimen for individual patients. In addition, the gene signatures and functional pathways inferred from the sequencing data in plasma would suggest potential targets to study the mechanisms of different responses to the treatment.

### Functional targets revealed by the multi-omics data

Our data have revealed many enriched pathways related to the tumor and its microenvironment (Fig. 4). For instance, one of the enriched pathways we found, focal adhesion, plays an essential role in cellular communication, which is also highly associated with cancer progress(33). As another example, significant down-regulation of translation pathway was revealed by our cfRNA data in the cancer patients’ plasma, which was also observed in tumor-educated platelet (TEP)(34). In addition, the suppressed immune signatures in plasma are also concordant with the fraction change of active immune cells in tumors revealed by previous studies(29). Because cancer patients responded differently to immune therapy(35), these pathways and signatures need to be further examined in different cancer subtypes.

### Limitations of this study

In this study, state-of-the-art technologies have enabled us to simultaneously investigate cfDNAs, cfDNAs’ methylation, small and total cfRNAs in a small amount of plasma (2-3 mL). However, some conclusions could be biased by a specific technology we used. For instance, we used MeDIP because it requires less plasma (1-1.5 mL) than bisulfite sequencing method (5-8 mL)(36). A bisulfite-based method like scWGBS (single cell Whole Genome Bisulfite Sequencing) would provide more precise information on DNA methylation than cfMeDIP while demanding very high sequencing depth (>30x) (37). As far as we know, this paper is one of the first studies investigating multi-omics data in liquid biopsy. However, we only focused on a few gastrointestinal cancer patients. Therefore, a comprehensive understanding of the cell-free molecules’ landscape in more cancer types and subtypes is needed. A large cohort study from multiple clinical centers is necessary for a robust predicting model in the real world.

### Perspective of multi-omics study in liquid biopsy

Like recent cfRNA studies(38, 39), we did not enrich EVs when sequencing cfRNAs, because the total cfRNAs in plasma include not only the EV-enriched cfRNAs but also those cfRNAs outside of the EVs (e.g., RNPs). More detailed studies of cfRNAs in different extracellular vehicles and RNPs would be useful. Moreover, the circulating system of human body is a highway for biological signal transporting by vesicle or in other forms, where cfDNA and cfRNA only represent part of the heavy traffic. Interpreting these biological processes in the circulating system requires more efforts to investigate more cell-free molecules such as proteins and lipids. Deciphering multiple cell-free molecules also needs many other technologies and experiments, such as cfChIP-seq(40).

## Materials and Methods

### Cohort design, sample collection and processing

We sequenced 360 cell-free omics datasets (86 cfWGS, 98 cfMeDIP-seq, 127 total cfRNA-seq, and 49 small cfRNA-seq) from 161 individuals (44 colorectal cancer patients, 36 stomach cancer patients, and 81 healthy donors) (Supplementary Table 1). Amon them, 95 were matched in 2-omics data, 84 were matched in 3-omics data, and 42 were matched in 4-omic data (Fig. 1a). By requiring enough coverage ratio and total mapped reads (see the detailed analyzing Methods below, Supplementary Tables 2,3), we kept most datasets (352 out of 360) for the downstream analyses.

The individuals were recruited from Peking University First Hospital (PKU, Beijing). Informed consent was obtained for all patients. Cell-free genome (cfWGS: individual number = 125), epigenome (cfMeDIP: individual number = 150), and transcriptome (total cfRNA-seq: individual number = 152; small cfRNA-seq: individual number = 73) were profiled. Patients’ age was distributed between 42-87 years (median age = 64 years), and most patients (51 out of 80) were diagnosed with stage I/II (Extended Data Fig. 1a). Different subtypes of colorectal and stomach cancer were included in the cohort (Extended Data Fig. 1b). For each person, 2-3 mL plasma sample was divided into 2-4 parts for 2 to 4-omics sequencing. For some samples (mostly from healthy donors), the plasma volumes were limited (less than 2 mL). We mixed these samples from persons with the same gender and similar age, then divided them into 2 to 4 parts (Extended Data Fig. 1c).

Peripheral whole blood samples were collected using EDTA-coated vacutainer tubes before any treatment of the patients. Plasma was separated within 2 hours after collection. All plasma samples were aliquoted and stored at −80°C before cfDNA and cfRNA extraction. Each sample was divided into 2-4 parts for sequencing different molecular types.

Peripheral blood mononuclear cells (PBMCs) were separated from whole blood by Ficoll (TBD, LTS1077-1). All PBMC samples were stored at −80°C.

Tissue samples were collected during surgery and transferred to liquid nitrogen within 30 minutes. Normal tissue adjacent tumor (NAT) was collected at least 2 cm away from the primary tumor.

### Isolation and sequencing of cfDNA (cfWGS) and cfDNA Methylation (cfMeDIP)

cfDNA was extracted from plasma using QIAamp MinElute ccfDNA Kit (Qiagen). DNA concentration was quantified by Qubit dsDNA HS Assay kit (Thermo Fisher Scientific). Up to 5 ng plasma cfDNA (~0.5 mL plasma) was used for cfWGS library with Kapa HiFi Hotstart ReadyMix (Roche) in 11-13 cycles. Libraries were sequenced on Illumina HiSeq X-ten (~60.7 million paired-end reads per library) with paired-end read length of 150 bases.

cfDNA methylation (cfMeDIP-seq) library was prepared following a previous protocol (41). Up to 15 ng plasma cfDNA (~1 mL plasma) were used as input, followed by end repair and A-tailing using Kapa Hyper Prep Kit (Kapa Biosystems). Next, adaptors were ligated using NEBNext Multiplex Oligos index (NEB). Phage lambda DNA was added to fill the low input to 100 ng. After heat-denature and snap-cool, single-stranded DNA mixture was incubated with 5-mC antibody provided by MagMeDIP-seq Package (Diagenode), followed by 14-16 cycles of library amplification, bead purification, and size selection. Libraries were sequenced on Illumina HiSeq X-ten (~42.9 million paired-end reads per library) with paired-end read length of 150 bases.

### cfWGS data processing and quality control

Raw fastq files were trimmed with *trim_galore* (All software being used in this study were summarized with versions and references in Supplementary Table 5.), then aligned to hg38(42) genome with default parameters using *bwa-mem2*. Reads were further filtered by proper template length (20 bp to 1000 bp) using *samtools* and de-duplicated using *GATK MarkDuplicates*. Base quality was recalibrated using *GATK BaseRecalibrator*.

We developed a set of quality control criteria to filter out poor libraries (Supplementary Table 2). 6 quality control steps were included: 1) relH score (the relative frequency of CpGs) < 1.5; 2) saturation score (300 bp bins correlation) > 0.9; 3) genome depth > 0.2; 4) coverage ratio > 0.1; 5) mapped ratio > 0.9; 6) unique read pairs > 2 million. Finally, 2 samples were filtered out.

### cfMeDIP-seq data processing and quality control

Methylation data were trimmed by *fastp*. Clean reads were firstly subjected to lambda genome alignment and then hg38(42) genome using *bowtie2* with “end-to-end” mode. Mapped reads were then de-duplicated by *GATK MarkDuplicates*. For the quality control procedure, we employed *MEDIPS* package to get CpG enrichment metrics and saturation estimation in 300 bp genome-wide bins. *featureCounts* were used to assign reads to each gene.

In data processing, we included 6 quality control steps (Supplementary Table 2): 1) saturation score > 0.9; 2) GoGe score (the observed/expected ratio of CpGs) > 1.2; 3) relH score > 1.5; 4) coverage ratio > 0.05; 5) mapped ratio> 0.9; 6) unique read pairs > 2 million. In total, 3 samples were filtered out.

### Isolation and sequencing of cfRNAs (total cfRNA-seq and small cfRNA-seq)

Total cfRNAs were extracted from ~1 mL of plasma using the Plasma/Serum Circulating RNA and Exosomal Purification kit (Norgen). Recombinant DNase I (TaKaRa) was used to digest DNAs. One set of ERCC RNA Spike-In Control Mixes (Ambion) was added. Next, the RNA Clean and Concentrator-5 kit (Zymo) was used to obtain pure total RNA. The total cfRNA library was prepared by SMARTer^®^ Stranded Total RNA-Seq Kit – Pico (TaKaRa). Libraries were sequenced on Illumina HiSeq X-ten (~37.5 million paired-end reads per library) with a length of 150 bases.

Small cfRNAs were extracted from ~1 mL of plasma using the miRNeasy Serum/Plasma Kit (Qiagen). 1ul ExiSEQ NGS Spike-in (Qiagen) was added to the extracted RNA. The small cfRNA library was prepared with the QIAseq miRNA Library Kit (Qiagen). Libraries were sequenced on Illumina HiSeq X-ten (~40.1 million reads per library), where adaptors linked to the short reads were later removed.

### Total cfRNA-seq data processing and quality control

For total cfRNA-seq data, adaptors and low-quality sequences were trimmed using *cutadapt*. Reads shorter than 16 nt were discarded. For template-switch-based RNA-seq data, GC oligos introduced in reverse transcription were trimmed off, after which reads shorter than 30 nt were discarded. The remaining reads were mapped to ERCC’s spike-in sequences, NCBI’s UniVec sequences (vector contamination), and human rRNA sequences sequentially using *STAR*. Then, all reads unmapped in previous steps were mapped to the hg38(42) genome index built with the GENCODE(43) v27 annotation. Reads unaligned to hg38 were aligned to circRNA junctions(44). For circRNA, only fragments spanning back-splicing junctions were taken into consideration. Duplicates in the aligned reads were removed using *GATK MarkDuplicates*. To avoid the impact of potential DNA contamination, only intron-spanning reads were considered for gene expression quantification(34). Intron-spanning reads were defined as a read pair with a CIGAR string in which at least one mate contains ‘N’ in the BAM files. Reads on exons were counted and aggerated to gene by *featureCounts*.

We filtered total cfRNA-seq samples using multiple quality control steps (Supplementary Table 3): 1) raw read pairs > 10 million; 2) clean read pairs (reads remained after trimming low quality and adaptor sequences) > 5 million; 3) aligned read pairs after duplicate removal (aligned to the hg38(42) human genome, and circRNA junctions) > 0.5 million; 4) fraction of spike-in read pairs < 0.5; 5) ratio of rRNA read pairs < 0.55; 6) ratio of mRNA and lncRNA read pairs > 0.2; 7) ratio of unclassified read pairs < 0.6; 8) number of intron-spanning read pairs > 100,000, 9) exonic/intronic reads ratio > 1. In total, 3 samples were filtered out.

### Small cfRNA-seq data processing and quality control

For small cfRNA-seq data, reads quality lower than 30 or length less than 15 were filtered by *trim_galore*. The remaining reads were sequentially mapped to ExiSEQ NGS Spike-in (a mix of 52 synthetic 5’ phosphorylated microRNAs), NCBI’s UniVec sequences, and human rRNA sequences, miRNA recorded in miRBase(45), lncRNA, mRNA, piRNA, snoRNA, snRNA, srpRNA, tRNA, transcripts of unknown potential (TUCPs) annotated in MiTranscriptome(46), Y_RNA by *bowtie2*. Mapped reads were sorted and indexed by *samtools*. Duplicates were removed by *umi_tools*.

We filtered small cfRNA-seq samples using 2 quality control steps (Supplementary Table 3): (1) datasets are required to have at least 100,000 reads that overlap with any annotated RNA transcript in the host genome, and (2) over 50% of the reads that map to the host genome also align to any RNA annotation. All small cfRNA-seq samples have enough reads for quantification, and most of the reads are aligned to RNA.

### Isolation and sequencing of RNAs in tissue cells and PBMCs

Tissue RNA was extracted by Trizol. The tissue RNA library was prepared with the NEBNext Ultra™ II RNA Library Prep Kit for Illumina. PBMC was seprated by Ficoll from whole blood. The PBMC RNA library was prepared by SMARTer^®^ Stranded Total RNA-Seq Kit – Pico (TaKaRa). All libraries were sequenced on Illumina HiSeq X-ten (~38.8 million per library) with paired-end read length of 150 bases, where adaptors being sequenced were later removed.

### Genome annotations

Human gene-centric genome regions and RNA biotypes were extracted from GENCODE v27 gtf file using *bedtools*. Human genome blacklist regions(47) were downloaded from ENCODE (https://www.encodeproject.org/). CpG island regions were downloaded from UCSC genome browser (http://genome.ucsc.edu/). CpG shore and shelf were defined as 2 kb and 4 kb flank regions, respectively. Repeated regions were downloaded from RepeatMasker (rmsk) database in UCSC genome browser. Promoter regions were defined as −2000 bp to +500 bp relative to TSS, according to a recent study(48).

### cfDNA and cfRNA length estimation

The length of cfDNA was summarized using BAM metric “TLEN” (Extended Data Fig. 2b). Insert length of total cfRNA-seq (Extended Data Fig. 2h) was estimated by MISO, using long constitutive exons as references.

### Correlation calculation among samples and omics

For correlation among samples, experiment reproducibility was checked using high throughput data correlation. Sample-based (i.e., sample A correlated with B by all genes abundances) Pearson correlations and corresponding *P*-values were calculated by *rcorr* function in R package *Hmisc*. Inter- or inner-omics correlations among different cancer types were averaged from multiple samples.

For gene correlation among omics, gene-based correlations (e.g., a gene’s DNA copy number and its RNA expression in the matched samples) were calculated. To compare omics correlation in cell lines and tissues, we downloaded RNA expression, DNA copy number, and DNA methylation data from the Cancer Cell Line Encyclopedia (CCLE)(20) and the Cancer Genome Atlas (TCGA)(21) from UCSC Xena (https://gdc.xenahubs.net/) and the Cancer Dependency Map portal (https://depmap.org/), respectively. Matched 3-omics data (33 stomach and 49 large intestine cell lines; 337 STAD, 307 COAD tissues) were selected for further analysis. For TCGA data, the gene-level copy number data were calculated by taking the segmental mean of the corresponding gene; the DNA methylation data were analyzed by calculating the CpG average beta value in the promoter region (2000 bp upstream and 500 bp downstream of TSS) of each gene; the gene expression data were converted to TPM (transcripts per million) data. Genes with NAs were removed.

### Calculation of multiple cfDNA variations

DNA copy number, window protection score (WPS), end-motif frequency, and fragment size were calculated based on the cfDNA-seq data. And DNA methylation of the promoter and CpG island was calculated based on cfMeDIP-seq data.

DNA copy number: Copy number was calculated as a gene-centric CPM (counts per million mapped regions) using cfWGS data, where hg38 blacklist regions(47) were masked. It was standardized as z-score using HDs’ distribution.

WPS: Windowed protection score (WPS) was calculated as the originally described study with minor modifications to estimate nucleosome occupancy in cfDNA(49). In brief, we used similar parameters as previously described: a minimum fragment size of 120 bp, a maximum fragment size of 180 bp, and a window of 120 bp. To account for variations in sequencing depth between samples, we performed a normalization step by dividing the WPS by the mean depth of randomly selected 1000 background sites in the genome. And then, for each gene, we quantify the nucleosome occupancy in TSS by computing the mean WPS from −150bp to +50bp around TSS.

End-motif frequency: End-motif was calculated following Jiang et al.(50). In short, the occurrence of all 5’ end 4-mer sequences (256 in total) of each valid template were counted and normalized as a ratio for each sample. Shannon entropy was calculated from the frequency of motif as motif diversity score (MDS) for each sample (theoretical scale: [0,1]).

Fragment size: The fragment size ratio matrix was calculated following Cristiano et al.(51). In short, 100-150 bp and 151-220 bp cfDNA templates were defined as short and long fragments respectively, 504 filtered bins mentioned in the original paper were converted to 469 bins in GRCh38 genome version, the read counts of each fragments type were also adjusted by LOESS-based GC content correction model.

DNA methylation: For each sample, raw counts of cfMeDIP-seq in promoter regions were normalized to CPM for cfDNA methylation level. We also computed counts per 300bp non-overlapping windows, normalized to CPM, and reduced to windows encompassing CpG islands, shores, and shelves.

### Calculation of multiple cfRNA variations

All the RNA variations, except for miRNA abundance, were calculated based on the total cfRNA-seq data.

RNA expression/cfRNA abundance: raw counts of miRNAs were normalized to CPM using small cfRNA-seq data; raw counts of the other genes were normalized to TPM using total cfRNA-seq data.

cfRNA alternative promoter: transcript isoform abundance was quantified by *salmon* and normalized to TPM. TPMs of isoforms with transcript start sites within 10 bp (sharing the same promoter) were aggregated as one promoter activity. TPM < 1 promoter is defined as an inactive promoter. The promoter with the highest relative promoter activity is defined as the major promoter. The remaining promoters are defined as minor promoters (52).

cfRNA SNV: intron-spanning reads were split by *GATK SplitNCigarReads* for confident SNP calling at RNA level. Then, alterations were identified by *GATK HaplotypeCaller* and filtered by *GATK VariantFilteration* with the following 4 criteria: strand bias defined by fisher exact test phred-scaled *P*-value (FS) < 20, variant confidence (QUAL) divided by the unfiltered depth (QD) > 2, total number of reads at the variant site (DP) > 10, SNP quality (QUAL) > 20. Allele fraction was defined as allele count divided by total count (reference count and allele count).

cfRNA editing: *GATK ASEReadCounter* was used to identify editing sites based on REDIportal(53). The editing ratio was defined as allele count divided by total count.

cfRNA allele specific expression: *GATK ASEReadCounter* were used to identify allele specific expression gene site based on SNP sites. For each individual, Allelic expression (AE, AE = |0.5 - Reference ratio |, Reference ratio = Reference reads/Total reads) was calculated for all sites with ≥16 reads(54).

cfRNA splicing: The percent spliced-in (PSI) score of each alternative splicing event was calculated using *rMATs-turbo*.

Chimeric cfRNA: Reads unaligned to genome were remapped to chimeric junctions by *STAR-fusion* to identify chimeric RNA. Chimera references were based on GTex(55) and ChimerDB-v3(56).

Microbial cfRNA abundance: Reads unaligned to genome were classified using kraken2 with its standard database to identify microbial cfRNA at genus level. Potential contaminations were filtered according to previous study(12). Counts at the genus level were also normalized by total genera counts.

### Calculation of differential alteration between cancer and healthy control

cfDNA copy number, promoter methylation, and CpG island methylation: exactTest implemented in *edgeR* were used between cancer patients and HDs. |log2FC| > 0.59 and *P*-value < 0.05 was used as the cutoff for defining significant differential alteration.

cfDNA end motif and fragment size: each differentially used motif or differential size fragment were identified by the Wilcoxon rank sum test for relatively end motif usage or fragment size. *P*-value < 0.05 was used as the cutoff.

cfDNA window protection score: each differentially protected gene was identified by the Wilcoxon rank sum test for window protection score. |delta window protection score| > 0.5, and *P*-value < 0.05 was used as cutoff.

RNA expression/cfRNA abundance and cf-miRNA abundance: differentially expressed genes were identified using the exactTest method in *edgeR*. |log2FC| > 0.59 and *P*-value < 0.05 was used as cutoff.

cfRNA alternative promoter usage, editing, and SNV: each differentially used promoter or the differentially mutated allelic site or editing site was defined by the Wilcoxon rank sum test for promoter usage or allele fraction. |delta allele fraction| > 0.2 and *P*-value < 0.05 was used as cutoff.

cfRNA allele specific expression: each differentially expressed allelic site was defined by the Wilcoxon rank sum test for AE. |delta AE| > 0.1 and *P*-value < 0.05 was used as cutoff.

cfRNA splicing: differential splicing events were identified by the likelihood ratio test implemented in *rMATs*. |delta PSI| >= 0.05 and *P*-value < 0.05 was used as cutoff.

Chimeric cfRNA: differential chimeric RNA events were defined by the fisher exact test between cancer patients and healthy donors. |delta frequency| > 0.1 and *P*-value < 0.05 was used as cutoff.

Microbial cfRNA abundance: each differential genus abundance was defined by the Wilcoxon rank sum test. |delta AE| > 0.1 and *P*-value < 0.05 was used as cutoff.

### Pathway enrichment analysis

For the above differential alterations, up-regulated and down-regulated genes in cancer were annotated by Kyoto Encyclopedia of Genes and Genomes (KEGG)(57). For cfRNA SNP, allele specific expression, and editing, the dysregulated sites’ coordinates were assigned to the gene using an R package, *biomaRt*. KEGG enrichment was calculated using *clusterProfiler*.

### Integrative pathway analysis of multi-omics

Integrative pathway analysis of multi-omics data (i.e., RNA expression, CNA, DNA methylation) was performed using *ActivePathways(58). P*-values were corrected for multiple testing using the Holm procedure, and 0.05 was set as the cutoff value for significance. And then, the enrichment map was visualized using the plugin enhancedGraphics in *Cytoscape(59*).

### Cell type signature score calculation

Cell type signature scores were deconvoluted from the plasma/tissue total RNA-seq data, using CIBERSORTx(60) with 1000 permutations. CIBERSORTx uses a reference panel of signature genes of different cell types and implements a support vector regression model to estimate the compositions of a mixture of different cell types’ RNAs. We used panels of tumor microenvironment (TME) cells(61) and LM22 panels of immune cells(28). We also used TIDE(62) and EPIC(27) methods to calculate scores of TME cells. The input to CIBERSORTx, TIDE, and EPIC is the TPM read count matrix of cfRNA abundance. When calculating the score of EPIC:CAFs for the TCGA cohort, the CAF gene list was re-defined using our cfRNA-seq data (significantly correlated with the stage), and the input gene abundance values were derived from the tissue RNA-seq data of TCGA.

### Software

All software being used in this study was summarized with versions and references in Supplementary Table 5.

## Acknowledgments

This work is supported by Tsinghua University Spring Breeze Fund (2021Z99CFY022), National Natural Science Foundation of China (81972798, 32170671,81902384), National Key Research and Development Plan of China (2019YFC1315700), National Science and Technology Major Project of China (2018ZX10723204, 2018ZX10302205), Tsinghua University Guoqiang Institute Grant (2021GQG1020), Tsinghua University Initiative Scientific Research Program of Precision Medicine (2022ZLA003), Bioinformatics Platform of National Center for Protein Sciences (Beijing) (2021-NCPSB-005). This study was also supported by Beijing Advanced Innovation Center for Structural Biology, Bio-Computing Platform of Tsinghua University Branch of China National Center for Protein Sciences, Interdisciplinary Clinical Research Project of Peking University First Hospital and the Capital Health Research and Development of Special, Open Research Fund Program of Beijing National Research Center for Information Science and Technology.

## Funding for open access charge

Tsinghua University Spring Breeze Fund [2021Z99CFY022].

## Ethics approval and consent to participate

This study was approved by the institutional review board of Peking University First Hospital (2018-15). Informed consent was obtained from all patients.

## Consent for publication

All authors have approved the manuscript and agree with the publication.

