## Extended Data Fig. for "Cell-free multi-omics analysis reveals tumor status-informative signatures in gastrointestinal cancer patients’ plasma"

### Extended Data Figures

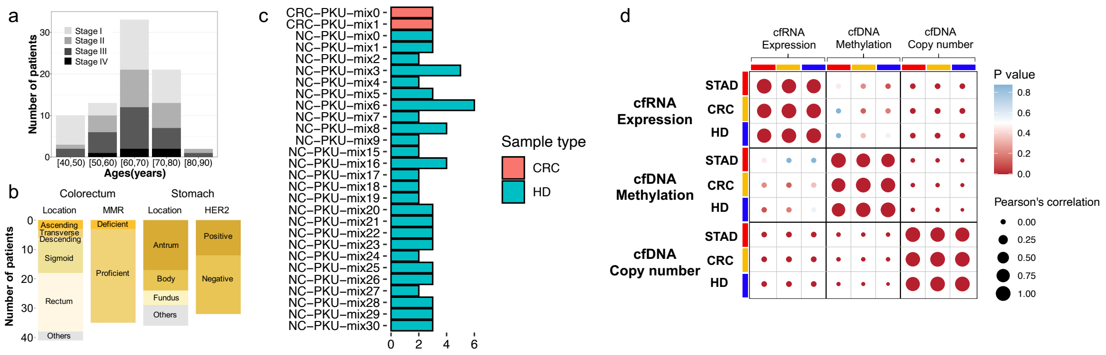

**Extended Data Figure 1. Clinical information and correlations of the multi-omics data**

**a,** Age and stage distribution of the patients. Age distribution of the patient cohort (range, 42–87 years; median, 64 years) highlighting disease stage.

**b,** Primary tumor location and molecular subtypes of the patients.

**c,** Illustration of the samples mixed by multiple individuals' plasma. Simultaneously sequencing cfDNA and cfRNA in the same individual consumes 2-3 ml plasma. In some cases, samples were mixed from individuals of same gender and similar age, due to the relatively small volume collected. More detailed usages of mixed samples are described in Supplementary Table 1.

**d,** Correlations of 77 samples with paired 3-omics data (cfWGS, cfMeDIP-seq, total cfRNA-seq).

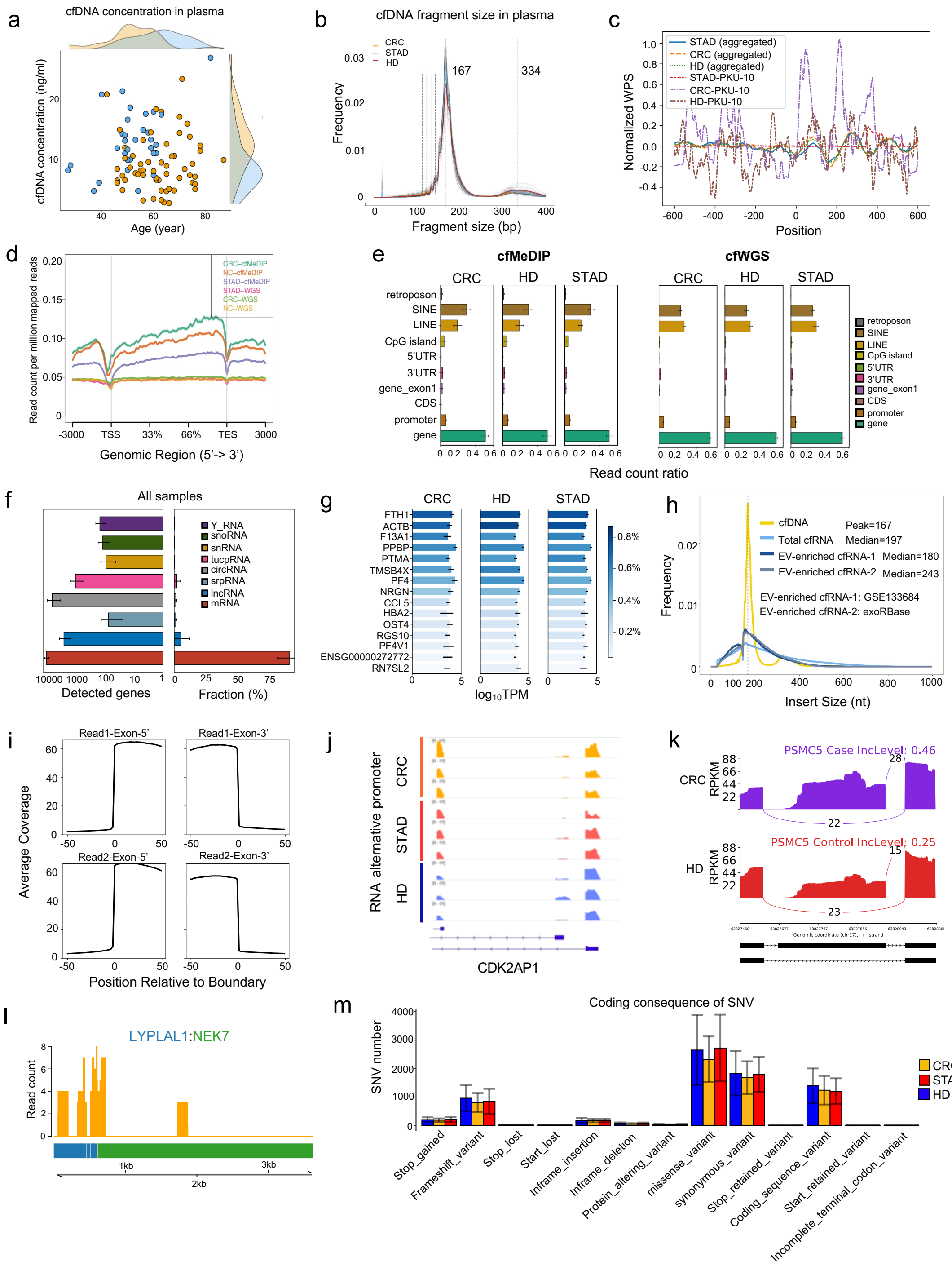

**Extended Data Figure 2. Basic characteristics of the cell-free multi- omics data.**

**a,** Concentration of the cfDNAs (ng/ml) in plasma for individuals with different age. Orange: tumor group, blue: normal group, side graphs show global distribution.

**b,** Distribution of the cfDNAs' fragment sizes in plasma. Average cfDNA fragment size in 3 different groups: colorectum cancer patients (yellow), stomach cancer patients (blude) and health donors (red). The most frequent fragment size is 167 bp and a second prevalent fragment size is 334bp. The fragment size is highly consistent with nucleosome protected DNA length.

**c,** Distribution of the cfDNA derived window protection scores (WPSs) around TSS.

**d,** Distribution of the cfWGS and cfMeDIP reads around TSS and TES. Y axis is RPM, x axis shows relative position to TSS and TES. RPM: read count per million, TSS: transcription start site, TES: transcription end site.

**e,** Distribution of the cfWGS and cfMeDIP reads in different genomic regions. Gene_exon1: representative first exon, Promoter: 2 kb upstream TSS to 0.5 kb downstream TSS, CDS: coding sequence, UTR: un-transcriptional region.

**f,** Detected gene number and fraction of reads for the total cfRNA-seq data. All RNA biotypes are annotated by GENCODE V27, except for tucpRNA, which is annotated by MiTranscriptome.

**g,** Top 15 genes ($\log_{10}$TPM TPM) sorted by the total cfRNA-seq reads. Color shows the fraction of total reads. Among the top expressed genes, we found housekeeping genes (ACTB/PTMA), platelet-related genes (F13A1/PPBP/PF4/PF4V1), erythrocyte-related genes (FTH1/HBA2) and exosomal RNAs, such as RN7SL2 (a signal recognition particle RNA).

**h,** Insert size for cfRNA, evRNA (extracellular vesicle) and cfDNA. The average insertion length of cfRNA is ~200 nt, which is different from cfDNA. Light blue: our total cfRNA-seq data. Dark blue: small evRNA (sEV-RNA-1) data downloaded from exoRBase version 1.0[1]. Grey: small evRNA (sEV-RNA-2) data downloaded from GSE133684. Yellow: our cfWGS data.

**i,** Aggregating coverage of the total cfRNA-seq reads at the exon-intron boundaries. The sharp edge suggests limited DNA contamination in the total cfRNA-seq data.

**j,** Example: total cfRNA-seq reads showing the promoter usage of CDK2AP1.

**k,** Example: total cfRNA-seq reads showing a spliced exon of PSMC5.

**l,** Example: total cfRNA reads showing the fusion of LYPLAL1:NEK7.

**m,** Coding consequences of the cfRNA-derived single nucleotide variations (SNVs).

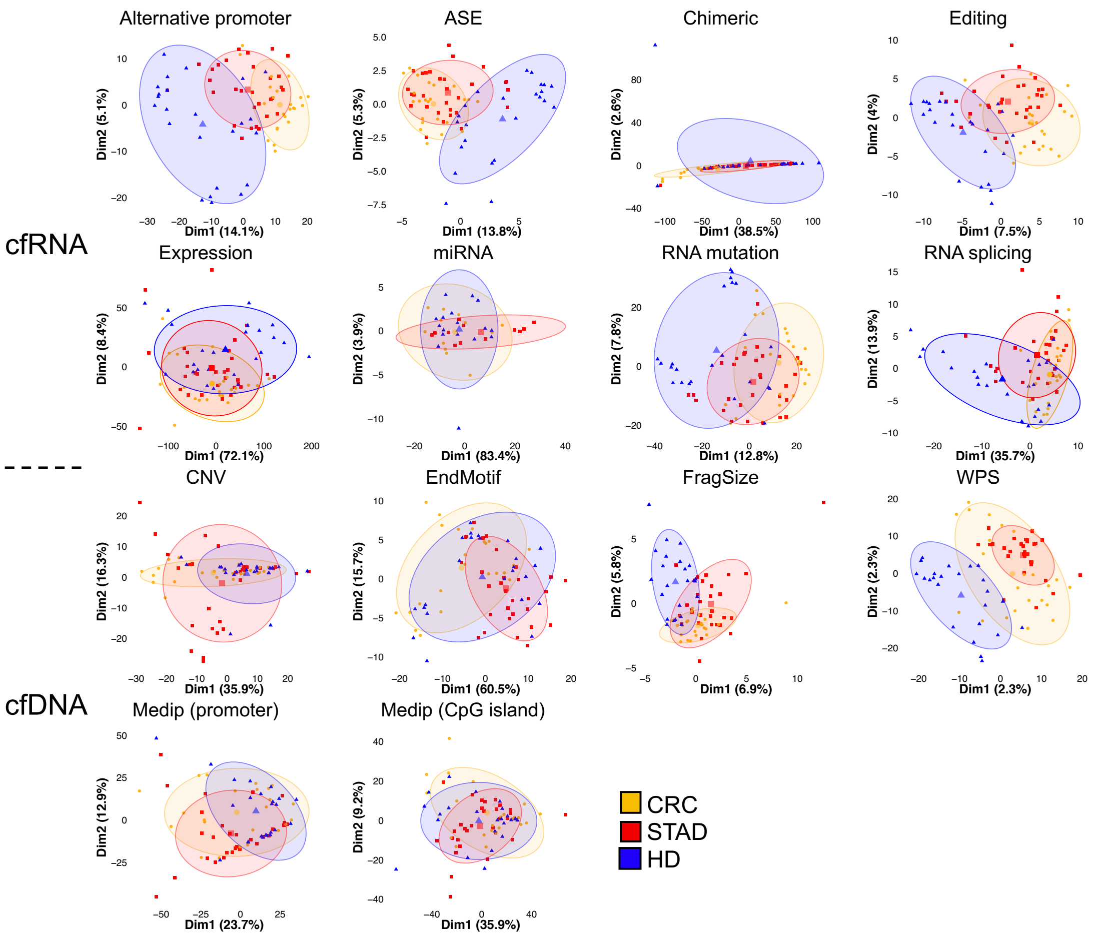

**Extended Data Figure 3. Principle component analysis of the differential alterations**

PCA analyses of all the differential alterations, including colorectal cancer (CRC) versus healthy donors (HDs), stomach cancer (STAD) versus HDs, and CRC versus STAD.

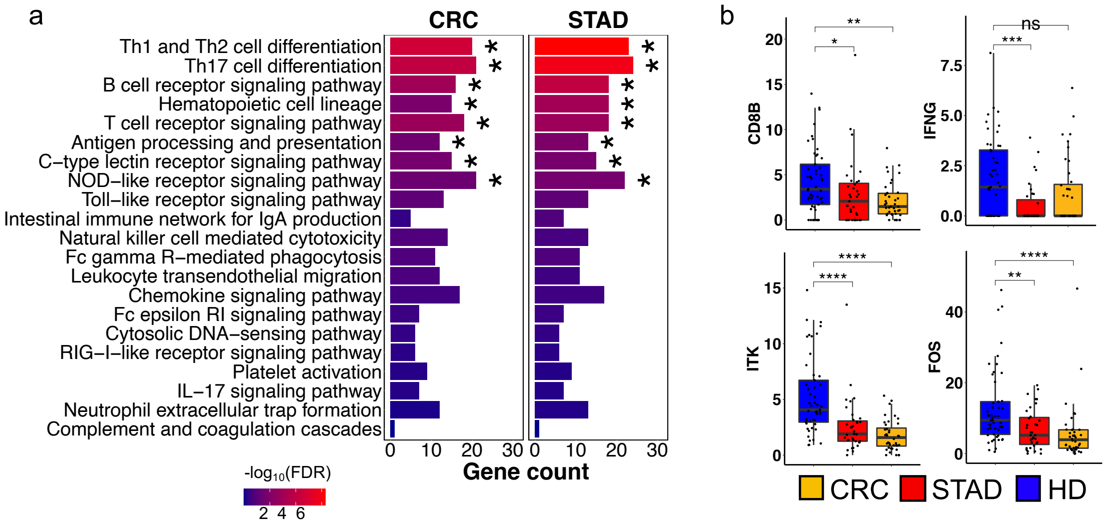

**Extended Data Figure 4. Downregulated immune pathways and genes defined by the total cfRNA-seq data.**

**a,** Immune related pathways enriched in CRC and STAD patients' plasma using the total cfRNA-seq data. *False Discovery Rate (FDR) < 0.05.

**b,** Example genes downregulated in the cancer patients' plasma. *CD8B:* a cell surface glycoprotein found on most cytotoxic T lymphocytes. *IFNG:* its primary producers are effector T cells and NK cells. *ITK:* caused a cascade transcriptional effect in stimulated T cells and diminished FOS (AP-1 gene family) in cancer patients[2].

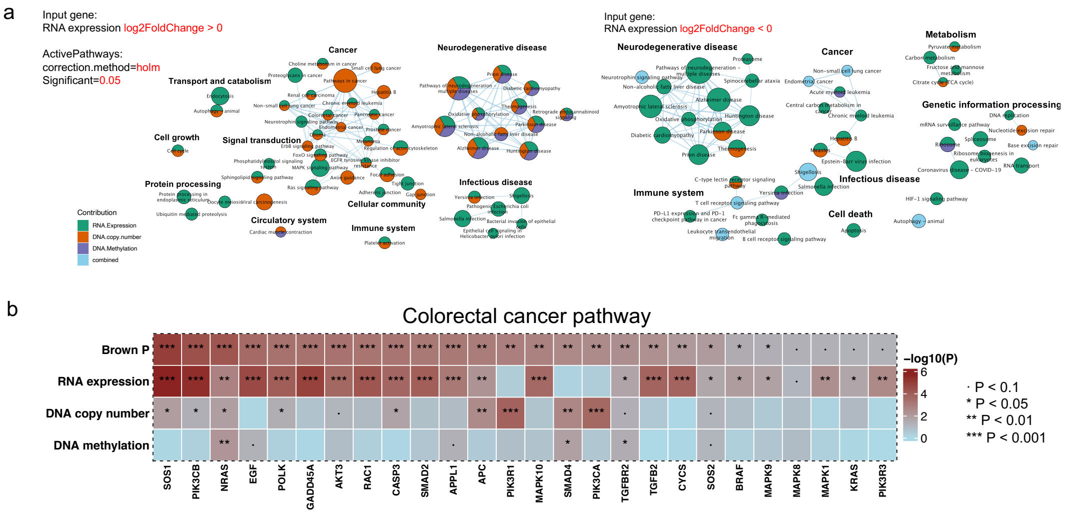

**Extended Data Figure 5. Multi-omics pathway enrichment analysis.**

**a,** Integrated networks of the enriched pathways. The networks were derived by *ActivePathways*[3] based on the multi-omics data. All circles' Holm's-method-corrected *P*-values are less than 0.05. Circle size represents gene number in each pathway; edge represents the similarity between each circle larger than 0.375; *P*-value of each gene represents significance of differential alteration.

**b,** Example genes altered at different omics in colorectal cancer pathway. Brown P was an integrated *P*-value of cfDNA copy number, cfDNA methylation (promoter) and cfRNA abundance by Brown's method. *P < 0.05, **P < 0.01, ***P < 0.001, ****P < 0.0001.

#### Extended Data References

1 Li, S. *et al.* exoRBase: a database of circRNA, lncRNA and mRNA in human blood exosomes. *Nucleic Acids Research* **46**, D106-D112 (2018).

2 Gallagher, M. P. *et al.* Hierarchy of signaling thresholds downstream of the T cell receptor and the Tec kinase ITK. *Proceedings of the National Academy of Sciences* **118**, e2025825118 (2021).

3 Paczkowska, M. *et al.* Integrative pathway enrichment analysis of multivariate omics data. *Nature Communications* **11**, 735 (2020).
